# ‘Optical Von-Frey’ method to determine nociceptive thresholds: a novel paradigm for preclinical pain assessment and analgesic screening

**DOI:** 10.1101/2023.11.02.565390

**Authors:** Jacqueline A Iredale, Amy J Pearl, Robert J Callister, Christopher V Dayas, Elizabeth E Manning, Brett A Graham

## Abstract

The goal of this study was to characterize a model that specifically activates peripheral nociceptors, allowing pure nociceptive thresholds to be monitored over a range of conditions including pathology or in screening analgesic compounds. Transgenic mice expressing channelrhodopsin-2 (ChR2) in cell populations positive for the transient receptor potential cation channel subfamily V member 1 (TRPV1) gene were bred to enable peripheral nociceptor photostimulation. Preliminary experiments confirmed the expected localisation pattern of ChR2 positive profiles in the dorsal root ganglion and superficial dorsal horn, mirroring TPRV1 expression. Brief hindpaw photostimulation with 470nm light caused hindpaw withdrawal and nocifensive behaviours in ChR2 positive animals but not control ChR2 negative animals. Using a simplified up/down approach, ‘optical’ nociceptive thresholds were assessed with a 5-intensity hindpaw photostimulation paradigm, establishing the minimum intensity required to produce a withdrawal response (optical threshold). All testing was also video recorded and analysed post-hoc to assess additional photostimulation evoked behaviours. Repeated testing over several days showed optical nociceptive thresholds and response duration were similar, supporting the stability of these variables across a timeframe relevant to onset of pathology or drug administration. Optical nociceptive thresholds were also assessed following morphine administration (30 mg/kg), which significantly raised thresholds, highlighting analgesic screening utility of this model. Together, these findings demonstrate the peripheral photostimulation with optical thresholding is a useful addition to the preclinical nociception assessment toolkit, with the key advantage of inducing a purely nociceptive response to a non-invasive, non-tissue damaging stimulus.

## INTRODUCTION

Despite decades of intensive preclinical research, progress to develop new pain treatments has remained slow and largely unsuccessful. While there are many potential explanations for this failure, questions have been raised about the validity and predictive capacity of current preclinical nociceptive assessment and analgesic screening tools, suggesting a need for novel approaches [10; 22; 32]. With no test able to directly measure pain in animals, assessments depend on evaluation of nociceptive responses, usually involving avoidance or nocifensive behaviours that are used as surrogate indicators of pain [4; 15; 28; 33]. The existing array of preclinical nociceptive assessment and analgesic screening tools include application of electrical, thermal, chemical, or mechanical stimuli to elicit a nociceptive response, commonly hindpaw withdrawal, licking, shaking, or grooming; and are typically used in conjunction with experimentally induced pain models that produce altered or pathological nociceptive signaling [4; 15; 28; 33].

In addition to characterising pain responses, tests can also specifically determine sensory thresholds, determining the minimum stimulus intensity required to provoke a hindpaw withdrawal response that can be reassessed during or following the onset of pathology or administration of experimental compounds [4]. Together, these approaches provide quantifiable measures related to nociception with clear face validity to the experience of pain. A key limitation of these thresholding-based approaches, however, is that stimuli generally activate a range of sensory receptor types (e.g. mechanoreceptive, thermoreceptive and chemoreceptive) [4; 15; 23]. Furthermore, high intensity stimuli are required to study nociception, carrying the risk of tissue damage. Fortunately, a range of technical and experimental advances have provided opportunities to develop new measures of pain and nociception in animals, expanding the preclinical assessment tools for studying pathological pain and assessing analgesic efficacy. For example, combinations of high-speed video imaging, statistical analysis, and machine learning are increasingly being employed to identify novel measures of spontaneous pain behaviour and automate this analysis [18].

Recent experimental advances have also provided new ways to activate neural pathways and study pain processing, including the fields of chemogenetics and optogenetics [16; 31; 35]. In the case of optogenetics, light sensitive proteins known as opsins can be specifically expressed in a target cell population, rendering these cells light-sensitive [27; 38]. Several studies have established that optogenetics can be used to activate peripheral nociceptors and study pain signalling evoked in the periphery, as well as activation at the level of the spinal cord [13; 19; 20; 35; 40]. While a range of strategies have been used to express channelrhodopsin-2 (ChR2) in nociceptive afferents and other cell populations, several studies have used the transient receptor potential cation channel subfamily V member 1 (TRPV1) to establish optogenetic control of nociceptors [5; 9; 12; 13; 21; 25; 26; 37].

Importantly, the TRPV1 ion channel mediates heat and chemically induced nociception, through expression in mechanically insensitive unmyelinated c-fibres [24; 30; 41]. Thus, photostimulation of this population in TRPV1::ChR2 mice, by applying 470 nm light to the central or peripheral nervous systems, can produce nociceptive signaling [5; 9; 12; 13; 21; 25; 26; 37]. Several studies have shown that under *in vivo* conditions peripheral optogenetic activation of nociceptors provokes nocifensive responses that can then be recorded and analysed [9; 30; 2; 13]. In building on this approach, work presented here describes an optical thresholding protocol, delivering set intensities of increasing photostimulation until a nocifensive hindpaw withdrawal response is elicited. This provides a form of nociceptive threshold that can then be monitored without tissue contact or injury and can assess changes in purely nociceptive signaling during the induction of pathological pain or as an analgesic efficacy screening model for novel compounds.

## METHODS

All experimental procedures were conducted at the University of Newcastle in accordance with the University’s Animal Care and Ethics Committee guidelines (protocol A-2021-134).

### Animals

Experimental mice were derived by crossing TRPV1^Cre^ with Ai32 (Jackson Laboratories, Bar Harbor, Maine, USA; lines #024109 and #017769, respectively). This generated offspring with ChR2/EYFP expressed in TRPV1 positive sensory neurons and their peripheral and central terminals. Both male and female mice (TRPV1^Cre^;Ai32; 3-6 months) were used as experimental animals. The uncrossed Ai32 parent line was used to confirm ChR2 activation underpinned responses to stimuli.

### Immunohistochemistry

Tissue was prepared for immunofluorescence confirmation of ChR2 expression and localization by animal overdose with pentobarbitone (800 mg/kg, i.p.; Virbac, NSW, Australia) followed by transcardial perfusion with 4% paraformaldehyde in phosphate buffer. The spinal cord and dorsal root ganglion (DRG) were there dissected free and post-fixed for an additional 2 hours in the same fixative. Transverse spinal cord sections prepared from the lumbar enlargement (L3-L5, 30 μm) embedded in TissueTek (Sakura Finetek, California, USA) and sectioned at -20°C on a Leica CM1950 cryostat (Leica, Wetzlar, Germany).

Alternatively, DRGs were polyethylene glycol (PEG) embedded after a series of washing and clearing steps: 3 x 15 min 80% ethanol (EtOH), 3 x 15 min dimethyl sulfoxide (DMSO) and then 3 x 15 min 100% EtOH. DRGs were then placed melted 1000MW PEG in a 46°C vacuum oven (Thermoline, NSW, Australia) until the tissue sank (approximately 2 hrs). The tissue was then embedded in 1450MW PEG, hardened, and sectioned (20 μm) on a microtome (American Optical, Illinois, USA).

Tissue sections from both the spinal cords and DRGs were placed in 10% normal donkey serum (Jackson ImmunoResearch, Pennsylvania, USA) in antibody diluent (0.3 M NaCl, 7.5 mM NaH2PO4, 0.3% TritonX-1000, 0.05% Na+ Azide), for 1 hour to block non-specific antibody binding. Sections were incubated overnight in primary antibodies against green-fluorescent protein (GFP, 1:1000 in antibody diluent, Abcam, ab13970), calcitonin gene-related peptide (CGRP, 1:200 in 0.1 M PBS, Abcam, ab36001) and neurofilament 200 (NF200, DRG sections only, 1:5000 in 0.1 M PBS, Sigma-Aldrich, N0142). After the primary incubation, sections were washed 3 x 15 min with 0.1 M PBS prior to a 2-hour incubation in secondary antibodies FITC (1:50 in 0.1 M PBS, Jackson ImmunoResearch, 703-095-155), Cy3 (1:50 in antibody diluent, Jackson ImmunoResearch, 715-165-147) and AMCA (DRG sections only, 1:50 in antibody diluent, Jackson ImmunoResearch, 715-155- 151) at room temperature. Sections were washed in 0.1 M PBS 3 x 15 min, mounted on slides with buffered glycerol and cover slipped. Slides were subsequently imaged on Olympus BX50 microscope (10X and 20X objectives), and analyzed using Image J software (NIH, New York, USA).

### In vivo photostimulation and behavioural testing

The testing apparatus consisted of a clear plexiglass cylinder was placed on top of an elevated plexiglass floor. A video camera was positioned in front of the testing setup to record all behaviours with 2 mirrors positioned behind the testing arena to capture behaviour from all angles. Under the elevated plexiglass floor, an LED driver (Thor Labs, New Jersey, USA) connected to 470 nm fibre-coupled LED (Thor Labs, New Jersey, USA) provided photostimulation through its attachment with a fibre optic patch cable, ceramic ferrule connection and optic fibre tip (300 μm) mounted on a fixed height stand. The LED driver was controlled by a Master-8 (AMPI, Jerusalem, Israel) pulse stimulator, delivering 1-second TTL pulses when triggered (Figure 1A). Set photostimulation intensities were calibrated using a digital optical power meter with a photodiode sensor positioned, sensor side down, on the elevated plexiglass flooring above the light path and the LED (Thor Labs, New Jersey, USA). This calibration procedure was performed on experimental days to ensure LED output accuracy.

**Figure 1.**
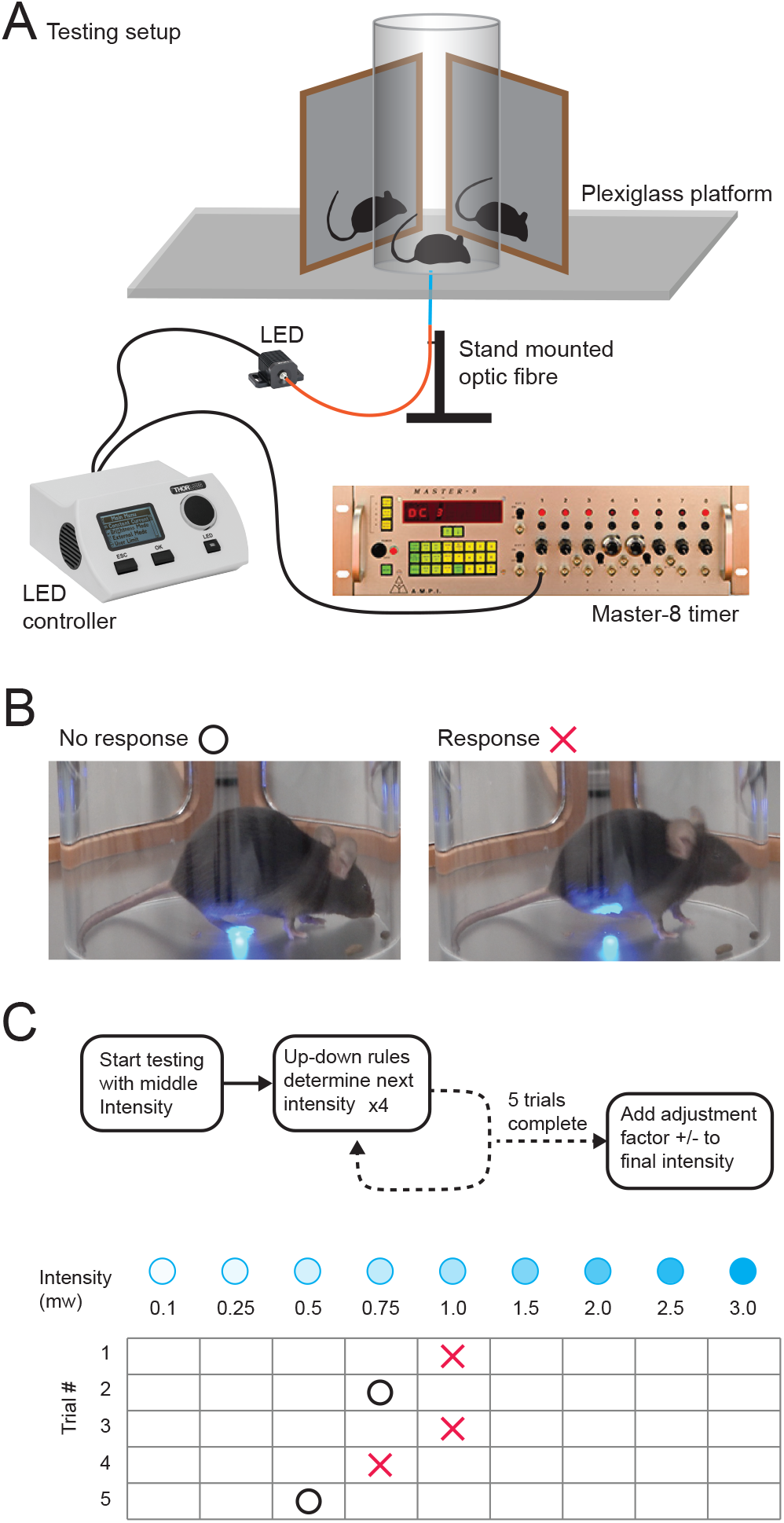
In vivo peripheral photostimulation experimental setup and procedure. (**A**) a TRPV1::ChR2 animal is placed in a clear plexiglass cylinder positioned on an elevated plexiglass platform. Two mirrors are positioned behind the cylinder so that the animal can be visualised from all angles with front on video camera recording. An LED driver is connected to 470 nm fibre-coupled LED, attached to a custom-mounted fibre optic patch cable under the elevated platform to deliver photostimulation through a 1 mm exposed fibre optic probe. The fibre optic probe tip is positioned a set distance from the underside of the elevated plexiglass floor to deliver consistent photostimulation intensities. The LED driver is triggered after positioning under the hindpaw of the mouse via a Master-8 pulse stimulator, which provides a timed 1 second TTL pulse. (**B**) images show hindpaw photostimulation with an examples of no response and response trials. **(C)** diagram summarizes 5-trial up/down paradigm for photostimulation. Following the first photostimulus at a set intensity of 1mW, subsequent photostimulus intensities are determined based on the result of the previous trial, increasing intensity if no withdrawal response to the prior intensity was observed, or decreasing if a withdrawal response was observed. A total of 5 photostimulation trials are conducted and the optical threshold assigned as the last photostimulation intensity +/- an adjustment factor of half the intensity to the next highest photostimulation intensity if there was no withdrawal response recorded in the last photostimulation trial, or half the intensity to the next lowest photostimulation intensity if the last photostimulation trial caused a withdrawal response.

All experiments were conducted under quiet, low-light conditions at 22 ± 1°C and 50% relative humidity. Animals were acclimatised in the testing arena for a minimum of 30 min or until responses to the novel environment were diminished (i.e. minimal rearing, pushing or scratching plexiglass, and unprovoked grooming). During acclimatization, the experimenter periodically adjusted the LED ferrule position under the floor, without triggering photostimulation, as would occur during testing. A minimum of three acclimatization sessions were performed prior to an experimental protocol commenced with at least 24 hrs between sessions. Following acclimatization, animals entered an experimental protocol. On the testing day, each animal was placed in the recording arena for 10 minutes prior to photostimulation, with video recording commenced to capture baseline activity. Following a simplified up-down approach modelled to that described by Bonin et al. [7] for von Frey filament testing, 5 photostimulation trials were applied, varying photostimulation intensity in an ‘up/down’ manner. The first stimulus trial was at the approximate photostimulation intensities (1mW) required to provoke a withdrawal response, determined in a pilot test cohort. The subsequent photostimulation intensity was determined by the up/down rules based on the response to the previous trial. Photostimulation intensity was increased if no response was recorded for the prior trial and decreased if the prior trial recorded a withdrawal response. Each trial commenced once the animal stopped moving and appeared to be settled, with the LED activated to deliver a 1-second light pulse to the right hindpaw and observing the response (Figure 1B). A three-minute time out between photostimulation trials allowed the animal behaviour to return to baseline. The response to the photostimulation (yes/no) was noted and the LED intensity was adjusted for the next photostimulation. All testing was scored in real time (response vs. no response) to determine the next photostimulation intensity value (Figure 1C). The optical threshold was assigned as the intensity of the last photostimulation intensity +/- an adjustment factor that was half the increment to the next intensity (up if there was no response or down if the last trial caused a response).

Testing trials were spaced 3-days apart, and three initial baseline optical threshold testing sessions were performed to test the stability of baseline optical thresholds and response behaviours (n=9). An additional fourth baseline test was undertaken in a subset of animals (n=6) at a later timepoint (14 days following conclusion of the initial baseline testing), which would exceed the time required to undertake analgesic screening. This was used to assess the long-term stability of optical thresholds and nociceptive responses for several repeat tests. A cohort of animals also underwent baseline optical threshold testing before being administered morphine (10 mg/kg, s.c.) and sham injection (isotonic saline) prior to additional optical threshold trials, also spaced 3-days apart.

Video recordings also captured each photostimulation trial for post hoc analysis using BORIS [17], an open-source event-logging software for video/audio coding and live observations.

Each file assessed for specific behaviours and conditions (Table 1). Analysis was performed frame-by-frame, with behaviours coded in the pre-photostimulation period (active or inactive), as well as during and post photostimulation (hindpaw lifting, tapping, grooming). Video recordings also determine the latency between photostimulation onset and the first behavioural response. Once coded, behaviour data (e.g., hindpaw lifting, tapping, or grooming) was exported in 1-second bins. Our analysis also grouped behaviours (e.g., nocifensive = lifting/tapping/grooming) and normalized these to account for photostimulation exposure when hindpaw withdrawal reduced the photostimulation period. This normalization multiplied the response duration by the fraction of the photostimulation period (1-second) that achieved hindpaw exposure.

**Table 1.**
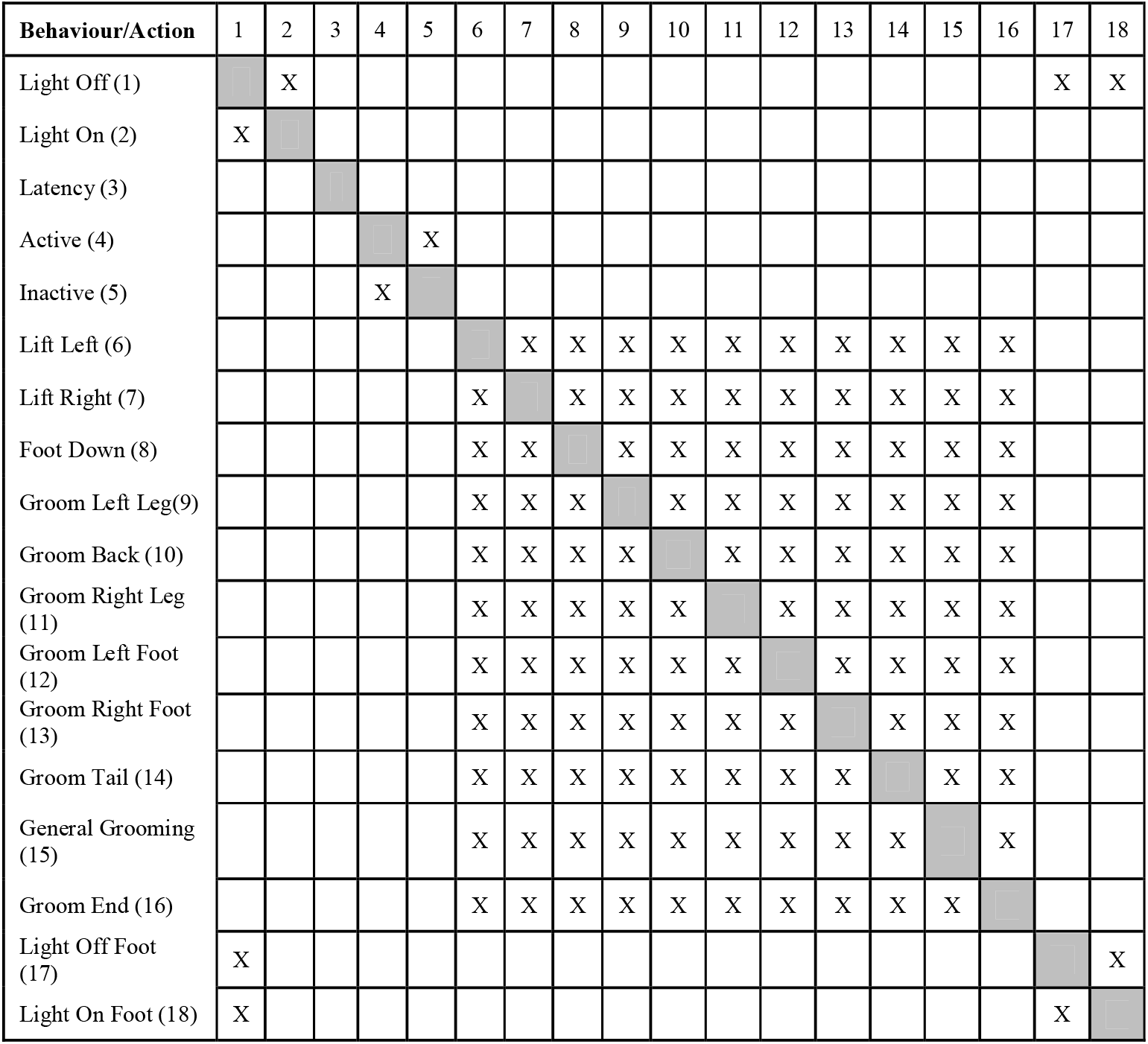
BORIS coded behaviours list and exclusion matrix for coded behaviours.

### Drugs and drug administration

Morphine (5 mg/mL; Pfizer, New york, USA) was prepared in isotonic saline was prepared immediately prior to administration. Morphine and vehicle (isotonic saline) were administered s.c. into the scruff of the neck at a volume of 5 mL/kg, and morphine at a dosage of 10 mg/kg.

### Statistics

Data were analyzed using the SPSS Statistical package (V28, IBM, New York, USA). Data were first examined to determine whether it satisfied the assumptions underlying analysis of variance. Repeated measures ANOVAs paired with post-hoc Tukey’s HSD or the non-parametric equivalent Friedman test paired with post-hoc Wilcoxon signed-rank tests were used to compare parametric and non-parametric data, respectively. All data are presented as mean ± standard error of the mean (SEM) and statistical significance was set at p ≤ 0.05.

## RESULTS

### Selective ChR2 expression and peripheral photostimulation responses

Several studies have validated the outcome of crossing of TRPV1^cre^ and cre-dependent ChR2 expressing transgenic mouse lines, however, the fidelity of ChR2-expression in animals used for these experiments and specificity of photostimulation responses was first assessed.

Immunolabeling analysis was undertaken in DRG and spinal cord tissue from a group of animals (n = 4). In DRG sections, ChR2/GFP (green) expression showed strong overlap with CGRP+ profiles (red) consistent with high expression in peptidergic nociceptors (Figure 2A- D). In contrast, there was minimal overlap in ChR2/GFP (green) expression with NF200+ profiles (blue), supporting negligible ChR2/GFP expression in large diameter myelinated afferents (Figure 2A-D). Group data comparisons confirmed that 51.33% ± 12.60% of ChR2+ cells co-expressed CGRP, whereas co-expression of ChR2 and NF200 was rare, accounting for only 3.43% ± 0.47% of ChR2+ cells (Figure 2E). Conversely, ChR2 expression was present in 65.22% ± 10.26% of the CGRP+ population but only 11.66% ± 1.00% of NF200+ profiles (Figure 2E). Comparisons of DRG cell diameter also showed that ChR2+ and CRGP+ populations were small diameter (TRPV1::ChR2: 19.49 ± 1.10 μm; CGRP: 21.33 ± 0.55 μm), unlike the larger diameter NF200+ DRG population (30.35 ± 1.15 μm) (Figure 2F). In the spinal cord, ChR2/GFP expression was restricted to the superficial dorsal horn, overlapping with CGRP labelling and consistent with the distribution of TRPV1+ terminals (Figure 2G-I) [5; 11; 37]. Together, these observations in DRG and spinal cord are consistent with selective expression of ChR2 in nociceptive, and not low threshold mechanosensitive afferents [13; 37].

**Figure 2.**
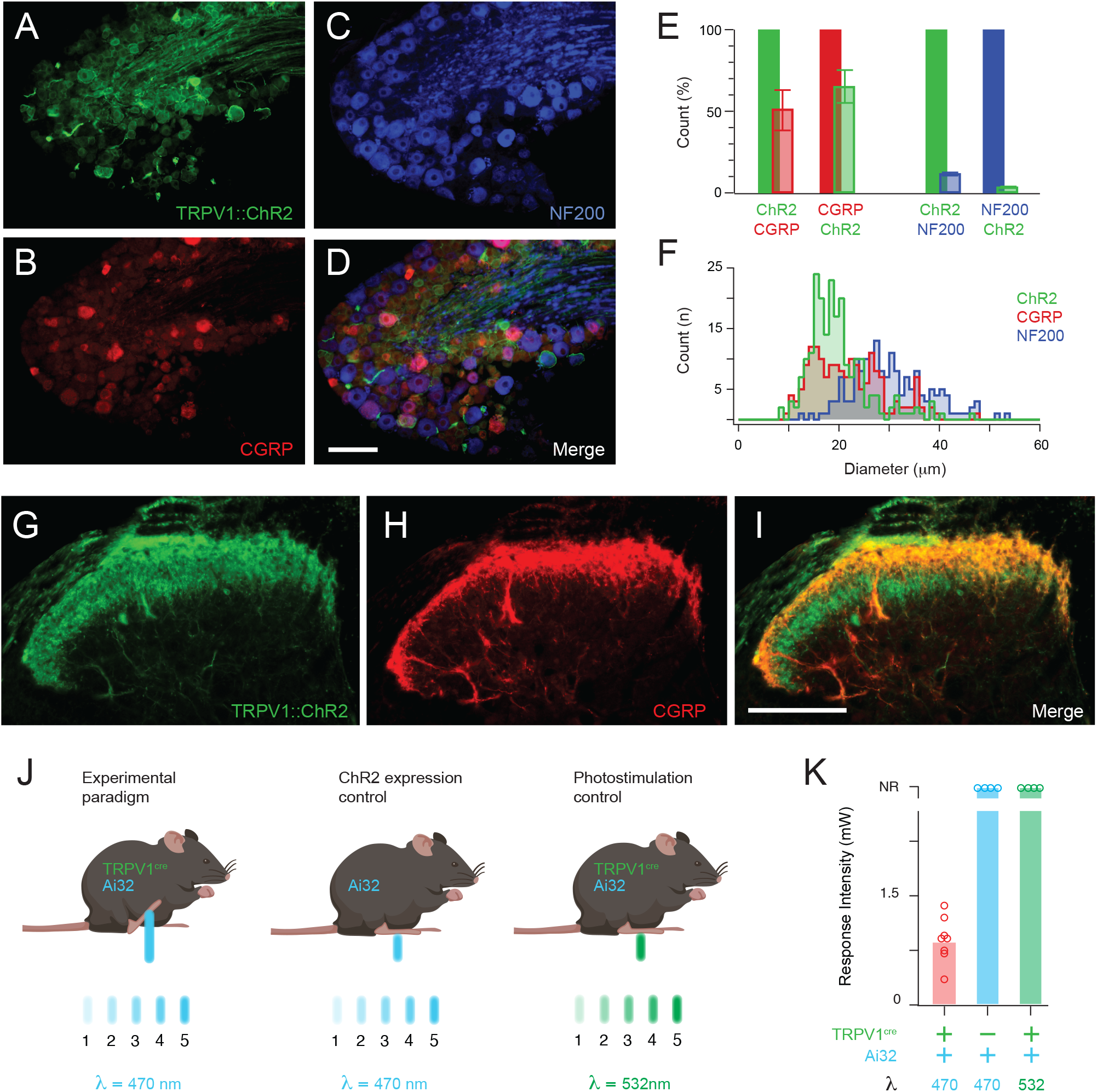
Expression of ChR2 in DRG and spinal dorsal horn of TRPV1::ChR2 mice and validation of selective photostimulation responses. **A-D,** images show distribution of ChR2/GFP (green, **A**), CGRP (red, **B**) and NF200 (blue, **C**) expression in DRG section, taken from a TRPV1::ChR2 mouse. Merged image (**D**) shows significant overlap of ChR2/GFP and CGRP-labelled cell profiles confirming the expected expression in small diameter peptidergic DRG cells. The lack of overlap between ChR2/GFP and NF200 confirms expression is largely excluded from large diameter myelinated DRG cells. Plots compare marker overlap (**E**) supporting significant overlap between ChR2/GFP and CGRP with minimal overlap between NF200 and either ChR2/GFP or CGRP. This relationship also correlates with the cell diameter for each population (**F**), with significant overlap between ChR2/GFP and CGRP, both having an average of approximately 20 μm, and minimal overlap between NF200 and either ChR2/GFP or CGRP, with NF200 averaging approximately 30 μm. **G-I**, images show a spinal dorsal horn (DH) section with ChR2/GFP (green, **G**) and CGRP (red, **H**) labelling. The merged immunolabelling (**I**) shows significant overlap of ChR2/GFP and CGRP profiles, consistent with ChR2/GFP expression limited to nociceptive afferents terminals. J, to confirm nocifensive responses were specific to ChR2 activation in the TRPV1::ChR2 transgenic animal line two control models were assessed. The first assessed the uncrossed parent line for the TRPV1::ChR2 animal, the Ai32 animal (n = 4), lacking TRPV1:cre to enable ChR2 expression. The second control used crossed TRPV1::ChR2 animals but instead of 470 nm blue light, photostimulation was applied at 532 nm light (green), outside the peak ChR2 excitation wavelength (n = 4). **K**, importantly, no response to peripheral photostimulation was observed in either control experiment at intensities up to 3 mW, exceeding the photostimulation intensity required to evoke responses in peripheral photostimulation of TRPV1::ChR2 mice (n=8). **K**, group data plots highlight the lack of response in either control model, confirming peripheral photostimulation are specific to TRPV1::ChR2 animals. All scale bars: 100 μm. Abbreviation NR = No response.

To confirm the selective ability of in vivo peripheral photostimulation in TRPV1::ChR2 animals (n = 8) to evoked nocifensive responses, two important control cohorts were also assessed (Figure 2J). One control included a subset of animals (n = 4) from the uncrossed Ai32 parent line where a loxP-flanked STOP cassette prevents ChR2 expression. The second control used crossed TRPV1::ChR2 mice, allowing for ChR2-expression, but a peripheral photostimulation wavelength (532 nm) outside the peak excitation wavelength for ChR2 (470 nm). Consistent with the previous literature, peripheral photostimulation of TRPV1::ChR2 mice at 470 nm of varying intensities produced distinct behavioural responses indicative of nociception with a mean optical threshold for responses of ∼ 1mW. In contrast, Ai32 mice did not exhibit any behavioural responses to peripheral photostimulation at 470 nm, even at the highest intensity tested (3.0 mW, Figure 2K). Likewise, TRPV1::ChR2 mice did not respond to peripheral photostimulation at 532 nm (Figure 2K). Together, these results confirm that selective activation of nociceptive afferents by peripheral photostimulation in TRPV1::ChR2 mice produced behavioural responses and excludes the potential for non-specific photostimulation responses caused by hindpaw illumination or visual cues, unrelated to ChR2 activation.

### Grouping nocifensive response domains

In characterizing peripheral photostimulation responses, three main nocifensive behaviours were commonly observed. These were tapping, lifting or grooming of the stimulated hindpaw. Tapping and lifting appeared milder forms of nocifensive response, while grooming, often maintained for an extended significant duration, was interpreted as a more substantial response to photostimulation. To determine the most relevant approach to quantify response data we first analysed a subset of animals (n=3) over 3 trials, assessing 10 individual tests for each animal and investigating individual response components (tapping, lifting and grooming). Lifting always occurred in response to photostimulation (30/30 tests) and was sometimes accompanied by tapping (6/30 tests) or grooming (4/30 tests, Figure 3).

**Figure 3.**
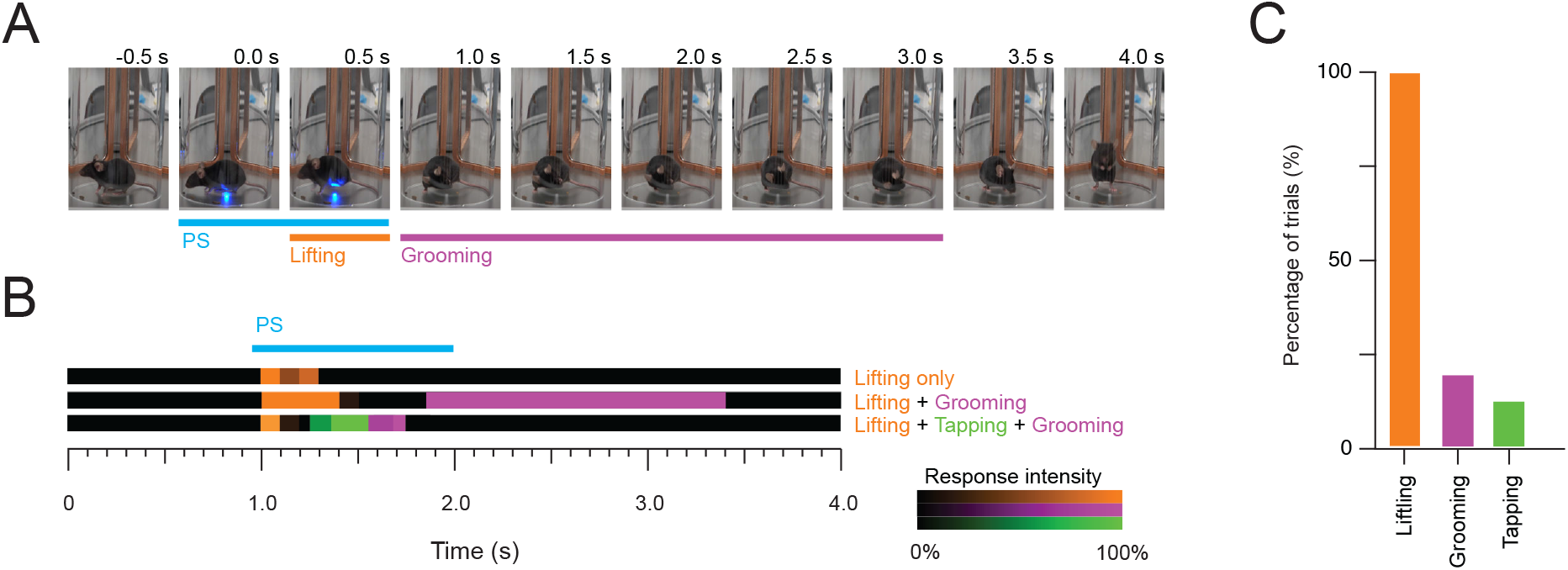
Classifying peripheral photostimulation evoked nocifensive responses. Initial assessment of photostimulation responses highlighted a range of different behaviours reflective of a nocifensive response including: lifting, tapping and grooming of the target hindpaw. **A**, images are consecutive stills from an example recording, showing the photostimulation period (blue line) frames including different behavioural responses (lifting- orange, grooming – purple). **B**, in order to determine the relative prevalence of each behaviour, video recordings were assessed from a subset of animals (n=3) over 3 trials, to determine the prevalence of each behaviour at threshold photostimulation intensity. Images shows analysis of 3 trails (different animals), with varying degrees of lifting, grooming and tapping. **C**, group data plots show paw lifting always occurred (30/30 trials) and was sometimes accompanied by tapping (6/30 trails) or grooming (7/30 trials). These observations supported grouping of the three behaviours for subsequent analyses collectively reflective the overall nocifensive response.

The contribution of tapping and grooming behaviours to the overall response often increased total response duration but never occurred in isolation from lifting, nor did they appear to reflect a significantly different response phenotype (Figure 3). Given that the consistent nature of paw lifting and variability of tapping and grooming, these beahviours were consequently grouped for subsequent nocifensive behaviour comparisons.

### Peripheral photostimulation repeatability and reliability

An important aspect of any viable assay assessing pathological pain or the efficacy of analgesic compounds is reproducibility over an anticipated testing period. Repeatability in the current peripheral photostimulation model was assessed in a group of TRPV1::ChR2 mice (n =10) tested across three trials, separated by 3-days. Comparisons assessed the optical threshold for paw withdrawal (lifting) during photostimulation, as well as withdrawal latency, and response duration. Group data comparisons confirmed that the optical threshold for withdrawal was stable across trials (p > 0.05), suggesting the utility of this measure to track changes in nociception (Figure 4). In contrast, withdrawal latency was more variable with the latency of trial 2 significantly shorter than trial 1 (0.45 ± 0.08 s vs. 0.85 ± 0.14 s, p < 0.05, Figure 4). Finally, despite variability in the response duration between animals and trials, mean values did not differ across trials (p > 0.05, Figure 4). Together, these results indicate some measures remain consistent over multiple testing days whereas others (eg, response latency) show greater variability. In a subset of experiments (n =8), animals underwent an additional photostimulation stability test 14 days after the third trial. Group data comparisons of the first 3 trials and the fourth trial showed that optical withdrawal threshold (0.87 ± 0.06 mW vs. 0.79 ± 0.22 mW), withdrawal latency (0.57 ± 0.06 s vs. 0.61 ± 0.14 s), and response duration (0.81 ± 0.23 s vs. 1.15 ± 0.52 s) did not differ over this period, further supporting the long-term stability of the peripheral photostimulation model (Figure 4).

**Figure 4.**
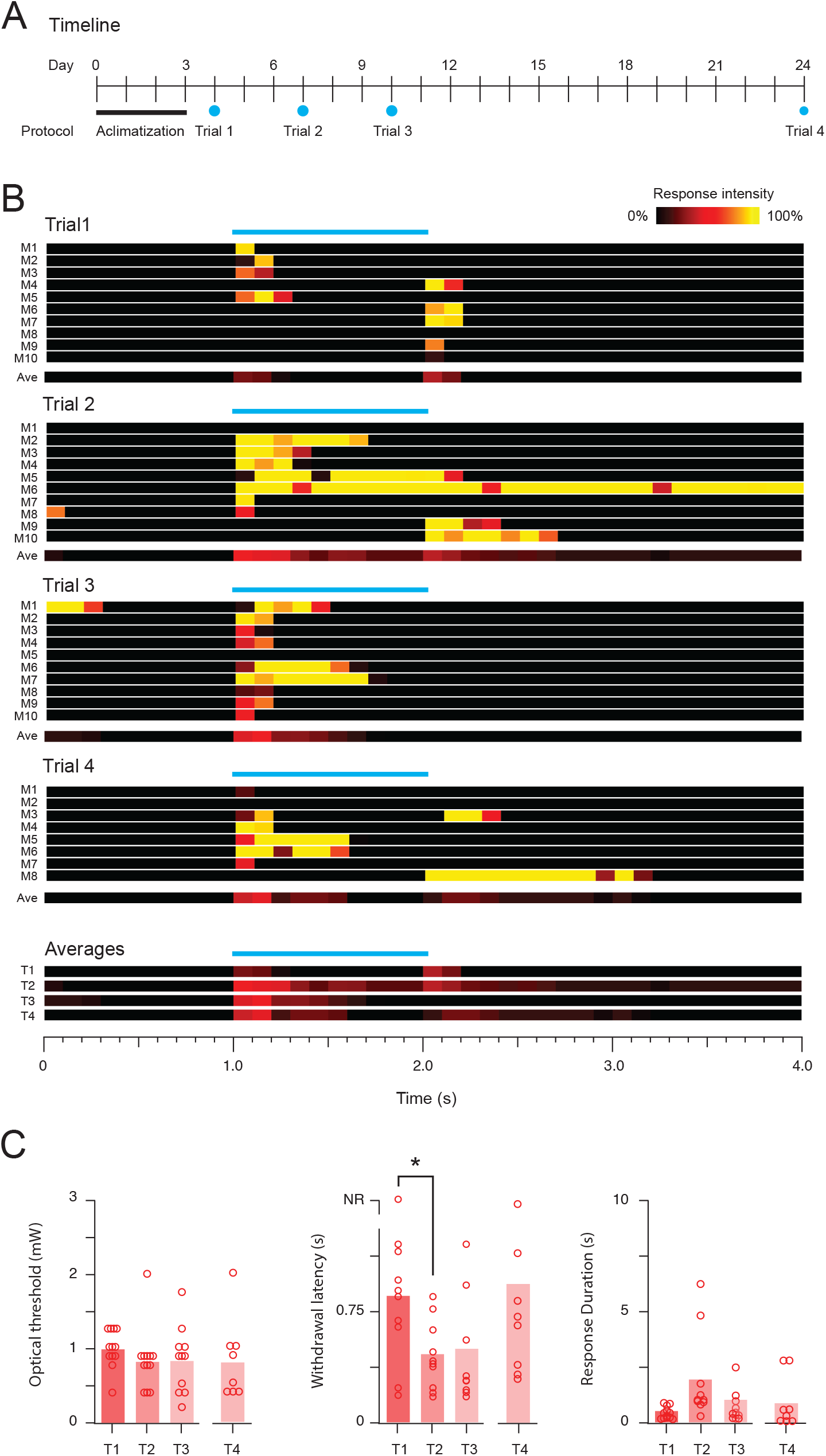
Peripheral photostimulation response stability testing. **A**, timeline summarizes peripheral phototstimulation testing schedule. All testing commenced with an acclimatisation period and three photostimulation trials were then performed with a 2-day break between trials, in some animals (n=8) a 4th trial was repeated, 14 days after the initial testing. **B,** panels show heatbar summaries of peripheral photostimulation responses across multiple trials. Nocifensive response intensity is denoted by colour and binned in 100 ms epochs. Each bar summarizes an individual mouses threshold photostimulation trail (M1-M10), with the group average presented below each trial number (1-4), also presented together for comparison (below). **C**, group data plots compare optical threshold, response latency, and response duration. Optical threshold was similar across all trials, whereas response latency differed between Trial 1 and Trial 2 (p < 0.05.). No other differences were seen across trials. Abbreviation NR = No response.

A necessary aspect of the up/down approach to optical withdrawal thresholding was the need to determine when withdrawal responses occurred as this determined the next photostimulation intensity. While full withdrawal of the hindpaw is a prominent movement that is easily detected, more subtle paw movements or adjustments related to photostimulation are potentially missed in real-time. Video recording of all trials made it possible assess each trial post hoc in slow motion to detect more subtle responses. This analysis confirmed that most withdrawal responses detected in real-time (79.84%, 293/367 assessed) accurately identified the onset of photostimulation associated behaviour. Of the remaining responses, slow-motion analysis detected photostimulation related movement of the hindpaw in 16.9% of trials that were originally classified as no response. This highlights even finer sensitivity may be possible if high speed videography and analysis were employed in the future. In addition, slow motion reanalysis identified a small number of photostimulation trials (3%), originally classified as containing a withdrawal response, that were reappraised as a spontaneous movement rather than a nocifensive response. Although this is a low error rate for small rodent behavioural analyses, and unlikely to influence the outcome of the up/down analysis, it also highlights scope for improvements that might be possible implementing computer vision and machine learning to remove human error.

### Analgesic sensitivity of peripheral photostimulation

To test the potential utility of peripheral photostimulation as an analgesic screening model, a cohort of animals was assessed before and after administration of a gold standard analgesic, morphine, as well as inclusion of a vehicle control saline injection (Figure 5, n = 7).

**Figure 5.**
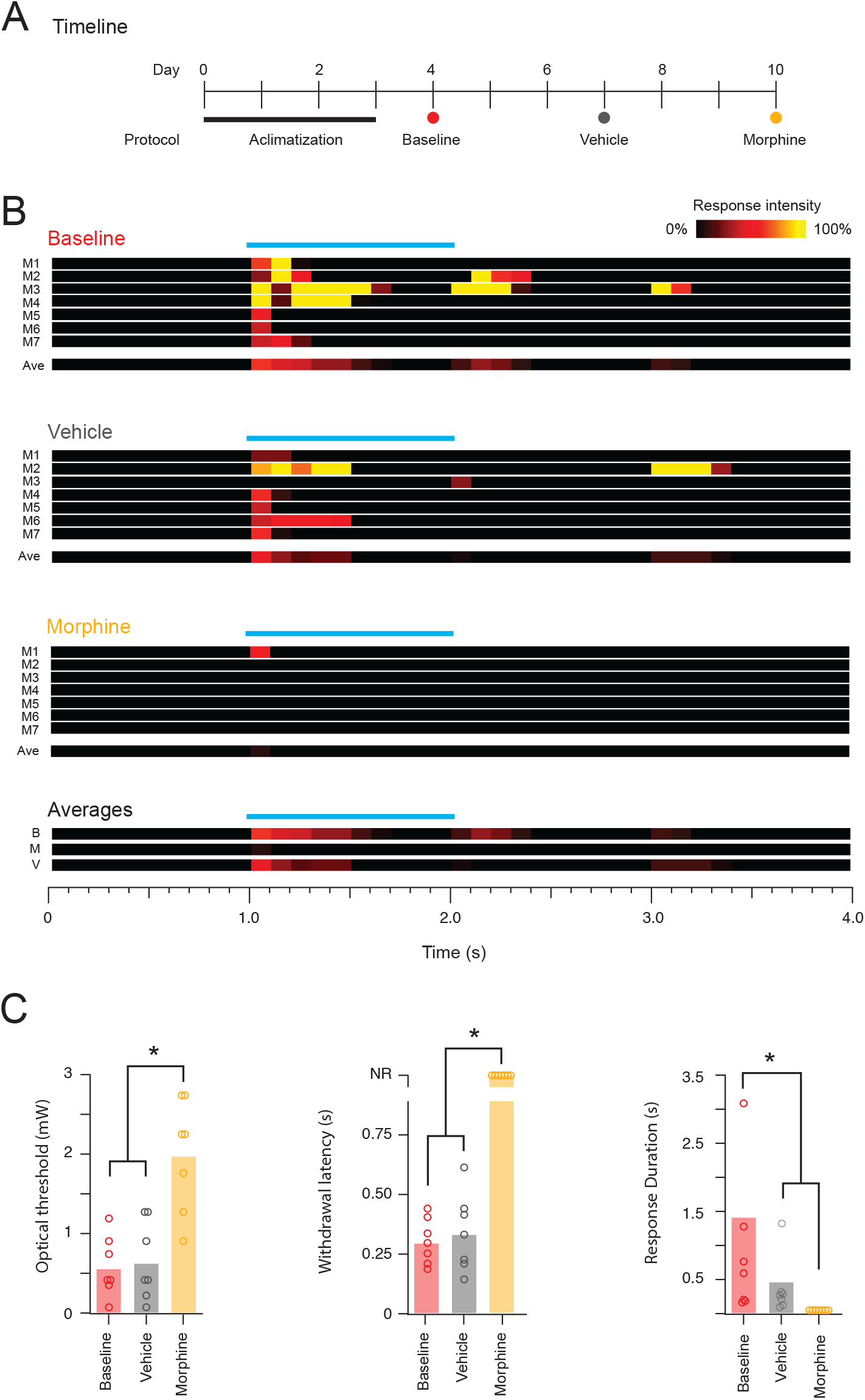
Effect of morphine on peripheral photostimulation response characteristics. **A**, timeline summarizes peripheral photostimulation testing schedule. All animals (n=7) underwent an initial acclimatization period before undergoing photostimulation trials at baseline, following vehicle injection (saline), and following morphine injection (30 mg/kg), with a 3-day break between trials. **B**, Panels show heatbar summaries of peripheral photostimulation responses in each mouse (M1-M7) across trials. Nocifensive response intensity is denoted by colour and binned in 100 ms epochs. Each heatbar shows an individual animals response at threshold photostimulation (determined in the baseline trial), with average for all animal below. C, group data plots compare optical threshold, response latency, and response duration. Following morphine administration (s.c.) optical threshold increased compared to both baseline and vehicle (p < 0.001). By extension, as the initial determined optical threshold did not provoke withdrawal responses and so response latency was assigned the maximum possible value of 1.5 s, and response duration the minimum value of 0. Both values differed significantly from both baseline and vehicle (p < 0.001, and p < 0.01, respectively). There were no significant differences detected between baseline and vehicle trials. Abbreviation NR = No response.

Importantly, no differences were detected in optical withdrawal thresholds between baseline and vehicle trials (0.55 ± 0.15 mW vs. 0.62 ± 0.19 mW), withdrawal latency (0.44 ± 0.06 s vs. 0.49 ± 0.09 s), and response duration (0.86 ± 0.40 s vs. 0.33 ± 0.16 s). In contrast, morphine significantly increased the optical withdrawal threshold compared to both baseline and vehicle (1.98 ± 0.27 mW vs. 0.55 ± 0.15 mW vs. 0.62 ± 0.19 mW, morphine vs. baseline vs. vehicle; p < 0.001; Figure 5). Given the change in optical threshold evoked by morphine, for threshold comparisons of withdrawal latency and duration, to baseline and vehicle results, morphine was assigned the maximum possible value of 1.5 s for latency and the minimum value of 0s for response duration. These results indicate the peripheral photostimulation model is sensitive to morphine analgesia supporting its utility as a novel analgesic efficacy screening tool.

## DISCUSSION

This study presents data supporting the use of optogenetics and peripheral photostimulation as a quantitative model to investigate nociception and assess the efficacy of potential analgesic compounds. We confirm that baseline photostimulation responses are stable across a time period that allows repeated trials for several days, with the optical withdrawal threshold the most reliable measure. Our data also confirms that the peripheral photostimulation model is sensitive to morphine, a prototypical analgesic, supporting the proposal that this model can provide an index of analgesic efficacy, also establishing reference data to benchmark other compounds against. Together, these findings suggest that addition of peripheral photostimulation to the preclinical nociceptive screening toolkit, using light rather than tissue damage and evoking a purely nociceptive signal, can enhance our ability to identify and develop new pain therapies.

It is important to recognise that several other studies have also utilized optogenetics to evoke peripheral photostimulation responses. Novel applications of this technology have continually grown since the first description of in vivo optogenetic nociceptive pathway activation, demonstrating photostimulation elicited acute behavioural responses, central sensitisation, and conditioned place aversion [13]. For example, precise control of peripheral afferents with photostimulation has allowed studies to characterise neural pathways and circuits [2; 3], demonstrate novel cytokine-mediated macrophage-nociceptor interactions [21], delineate an activity dependent protective molecular pathway for transient nociceptor sensitization [37] and resolve the role of specific afferents on sensory experience and pathology [34; 39].

Interestingly, while some studies employed the same TRPV1::ChR2 used here, many used different transgenic animals selectively targeting a range of primary afferents types including Ab fibres [39], tachykinin-1 (TAC1)-expressing [3], neuropeptide Y receptor-2 (NPYR2)- expressing [2], mas1-related G-protein-coupled receptor A3 (MrgprA3)-expressing [34], mast cell function-associated antigen (MAFA)-expressing [2], voltage-gated sodium channel Nav1.8 [13], and advillin-positive [29]. Applying the optical thresholding and compound screening approach described here across this range of animals represents a valuable opportunity for future drug discovery, mechanism, and efficacy studies. Likewise, the field of visceral nociception and pain research has incorporated optogenetic research approaches to selectively activate specific organ afferents using implantable LED systems [36]. Optical thresholding principles may also be applied here to rapidly advance drug discovery for visceral pain although a reliable and consistent behavioural readout would be required.

Consistent with the rapidly evolving use of optogenetics in pain research, our experiments also stand to benefit from a variety of developments. For example, a number of studies have used high-speed videography to improve the accuracy of response latencies recorded in comparable photostimulation paradigms [1; 2; 6; 8]. Implementation high-speed videography with optical threshold testing would likely provide additional insights on analgesic responses, although the simpler, more cost and time effective approach taken here was adequate to resolve a substantial morphine effect. Specific to the context of analgesic testing and discovery, the most effective compounds should produce large changes in sensitivity that would correspond to potent analgesia. Interestingly, recent description of an automated approach to photostimulation delivery and real-time analysis of behavioural responses offers an alternative approach to high-speed videography and would be amenable to implementation of optical thresholding [14]. This innovation also offers the opportunity for testing at greater scale, moving the optical thresholding approach toward a higher throughput screen.

One limitation of all nociceptive assessments based on reflex behaviour is their potential lack of relevance to the higher order experience and perception of pain. This issue has been elegantly addressed, combining optogenetics and a sensory detection task that allows animals to self-report detection of peripheral photostimulation in a lick-response paradigm [5]. Whilst more elaborate in the need for pre-training, this nociception self-report assay could also be combined with an optical thresholding procedure to track changes in nociceptive perception and analgesic screening. Likewise, Zhang et al. [42] have described an automated paw force, body and face grooming detection system, based on frustrated internal reflectance, for a simplified assessment of complex pain behaviours. Combined with the up/down optical thresholding paradigm described here, these innovations could constitute a multimodal readout of peripheral photostimulation to objectively screen analgesic efficacy and further enhance preclinical drug discovery.

## ACKNOWLEDGMENTS

This work was funded by the National Health and Medical Research Council (NHMRC) of Australia (grants 1144638, 1184974, and 2020113 to B.A.G.) and the Hunter Medical Research Institute (grant to B.A.G.).

## REFERENCES

[1] Abdus-Saboor I, Fried NT, Lay M, Burdge J, Swanson K, Fischer R, Jones J, Dong P, Cai W, Guo X. Development of a mouse pain scale using sub-second behavioral mapping and statistical modeling. Cell reports 2019;28(6):1623–1634. e1624.

[2] Arcourt A, Gorham L, Dhandapani R, Prato V, Taberner FJ, Wende H, Gangadharan V, Birchmeier C, Heppenstall PA, Lechner SG. Touch Receptor-Derived Sensory Information Alleviates Acute Pain Signaling and Fine-Tunes Nociceptive Reflex Coordination. Neuron 2017;93(1):179–193.

[3] Barik A, Thompson JH, Seltzer M, Ghitani N, Chesler AT. A brainstem-spinal circuit controlling nocifensive behavior. Neuron 2018;100(6):1491–1503. e1493.

[4] Barrot M. Tests and models of nociception and pain in rodents. Neuroscience 2012;211:39–50.

[5] Black CJ, Allawala AB, Bloye K, Vanent KN, Edhi MM, Saab CY, Borton DA. Automated and rapid self-report of nociception in transgenic mice. Scientific reports 2020;10(1):1–10.

[6] Blivis D, Haspel G, Mannes PZ, O’Donovan MJ, Iadarola MJ. Identification of a novel spinal nociceptive-motor gate control for Aδ pain stimuli in rats. Elife 2017;6:e23584.

[7] Bonin RP, Bories C, De Koninck Y. A simplified up-down method (SUDO) for measuring mechanical nociception in rodents using von Frey filaments. Molecular pain 2014;10(1):1–11.

[8] Browne LE, Latremoliere A, Lehnert BP, Grantham A, Ward C, Alexandre C, Costigan M, Michoud F, Roberson DP, Ginty DD. Time-resolved fast mammalian behavior reveals the complexity of protective pain responses. Cell reports 2017;20(1):89–98.

[9] Browne LE, Latremoliere A, Lehnert BP, Grantham A, Ward C, Alexandre C, Costigan M, Michoud F, Roberson DP, Ginty DD, Woolf CJ. Time-Resolved Fast Mammalian Behavior Reveals the Complexity of Protective Pain Responses. Cell Rep 2017;20(1):89–98.

[10] Burgess G, Williams D. The discovery and development of analgesics: new mechanisms, new modalities. J Clin Invest 2010;120(11):3753–3759.

[11] Cavanaugh DJ, Chesler AT, Bráz JM, Shah NM, Julius D, Basbaum AI. Restriction of transient receptor potential vanilloid-1 to the peptidergic subset of primary afferent neurons follows its developmental downregulation in nonpeptidergic neurons. J Neurosci 2011;31(28):10119–10127.

[12] Cohen JA, Edwards TN, Liu AW, Hirai T, Jones MR, Wu J, Li Y, Zhang S, Ho J, Davis BM, Albers KM, Kaplan DH. Cutaneous TRPV1+ Neurons Trigger Protective Innate Type 17 Anticipatory Immunity. Cell 2019;178(4):919–932.e914.

[13] Daou I, Tuttle AH, Longo G, Wieskopf JS, Bonin RP, Ase AR, Wood JN, De Koninck Y, Ribeiro-da-Silva A, Mogil JS, Séguéla P. Remote optogenetic activation and sensitization of pain pathways in freely moving mice. J Neurosci 2013;33(47):18631–18640.

[14] Dedek C, Azadgoleh MA, Prescott SA. Reproducible and fully automated testing of nocifensive behavior in mice. bioRxiv 2023:2023.2004. 2013.536768.

[15] Deuis JR, Dvorakova LS, Vetter I. Methods Used to Evaluate Pain Behaviors in Rodents. Front Mol Neurosci 2017;10:284.

[16] Duan B, Cheng L, Bourane S, Britz O, Padilla C, Garcia-Campmany L, Krashes M, Knowlton W, Velasquez T, Ren X. Identification of spinal circuits transmitting and gating mechanical pain. Cell 2014;159(6):1417–1432.

[17] Friard O, Gamba M. BORIS: a free, versatile open[source event logging software for video/audio coding and live observations. Methods in ecology and evolution 2016;7(11):1325–1330.

[18] Fried NT, Chamessian A, Zylka MJ, Abdus-Saboor I. Improving pain assessment in mice and rats with advanced videography and computational approaches. Pain 2020;161(7):1420–1424.

[19] Grajales-Reyes JG, Copits BA, Lie F, Yu Y, Avila R, Vogt SK, Huang Y, Banks AR, Rogers JA, Gereau IV RW. Surgical implantation of wireless, battery-free optoelectronic epidural implants for optogenetic manipulation of spinal cord circuits in mice. Nature protocols 2021;16(6):3072–3088.

[20] Iyer SM, Montgomery KL, Towne C, Lee SY, Ramakrishnan C, Deisseroth K, Delp SL. Virally mediated optogenetic excitation and inhibition of pain in freely moving nontransgenic mice. Nature Biotechnology 2014;32(3):274–278.

[21] Ji J, He Q, Luo X, Bang S, Matsuoka Y, McGinnis A, Nackley AG, Ji R-R. IL-23 enhances C-fiber-mediated and blue light-induced spontaneous pain in female mice. Frontiers in Immunology 2021;12.

[22] Kissin I. The development of new analgesics over the past 50 years: a lack of real breakthrough drugs. Anesth Analg 2010;110(3):780–789.

[23] Landerholm ÅH, Hansson PT. The perception threshold counterpart to dynamic and static mechanical allodynia assessed using von Frey filaments in peripheral neuropathic pain patients. Scandinavian Journal of Pain 2011;2(1):9–16.

[24] Lawson JJ, McIlwrath SL, Woodbury CJ, Davis BM, Koerber HR. TRPV1 unlike TRPV2 is restricted to a subset of mechanically insensitive cutaneous nociceptors responding to heat. J Pain 2008;9(4):298–308.

[25] Li J, Ali MSS, Lemon CH. TRPV1-Lineage Somatosensory Fibers Communicate with Taste Neurons in the Mouse Parabrachial Nucleus. The Journal of Neuroscience 2022;42(9):1719.

[26] Michoud F, Seehus C, Schönle P, Brun N, Taub D, Zhang Z, Jain A, Furfaro I, Akouissi O, Moon R, Meier P, Galan K, Doyle B, Tetreault M, Talbot S, Browne LE, Huang Q, Woolf CJ, Lacour SP. Epineural optogenetic activation of nociceptors initiates and amplifies inflammation. Nature biotechnology 2021;39(2):179–185.

[27] Mickle AD, Gereau RWt. A bright future? Optogenetics in the periphery for pain research and therapy. Pain 2018;159 Suppl 1(Suppl 1):S65-s73.

[28] Mogil JS. Animal models of pain: progress and challenges. Nature Reviews Neuroscience 2009;10(4):283–294.

[29] Park SI, Brenner DS, Shin G, Morgan CD, Copits BA, Chung HU, Pullen MY, Noh KN, Davidson S, Oh SJ, Yoon J, Jang KI, Samineni VK, Norman M, Grajales-Reyes JG, Vogt SK, Sundaram SS, Wilson KM, Ha JS, Xu R, Pan T, Kim TI, Huang Y, Montana MC, Golden JP, Bruchas MR, Gereau RWt, Rogers JA. Soft, stretchable, fully implantable miniaturized optoelectronic systems for wireless optogenetics. Nat Biotechnol 2015;33(12):1280–1286.

[30] Peirs C, Seal RP. Neural circuits for pain: recent advances and current views. Science 2016;354(6312):578-584.

[31] Peirs C, Williams S-PG, Zhao X, Arokiaraj CM, Ferreira DW, Noh M-c, Smith KM, Halder P, Corrigan KA, Gedeon JY. Mechanical allodynia circuitry in the dorsal horn is defined by the nature of the injury. Neuron 2021;109(1):73–90. e77.

[32] Percie du Sert N, Rice AS. Improving the translation of analgesic drugs to the clinic: animal models of neuropathic pain. Br J Pharmacol 2014;171(12):2951–2963.

[33] Sandkuhler J. Models and mechanisms of hyperalgesia and allodynia. Physiological reviews 2009;89(2):707–758.

[34] Sharif B, Ase AR, Ribeiro-da-Silva A, Séguéla P. Differential coding of itch and pain by a subpopulation of primary afferent neurons. Neuron 2020;106(6):940–951. e944.

[35] Smith KM, Browne TJ, Davis OC, Coyle A, Boyle KA, Watanabe M, Dickinson SA, Iredale JA, Gradwell MA, Jobling P. Calretinin positive neurons form an excitatory amplifier network in the spinal cord dorsal horn. Elife 2019;8.

[36] Spencer NJ, Hu H. Enteric nervous system: sensory transduction, neural circuits and gastrointestinal motility. Nature reviews Gastroenterology & hepatology 2020;17(6):338–351.

[37] Stemkowski P, García-Caballero A, Gadotti VDM, M’Dahoma S, Huang S, Black SAG, Chen L, Souza IA, Zhang Z, Zamponi GW. TRPV1 nociceptor activity initiates USP5/T-type channel-mediated plasticity. Cell reports 2016;17(11):2901–2912.

[38] Tan P, He L, Huang Y, Zhou Y. Optophysiology: Illuminating cell physiology with optogenetics. Physiological Reviews 2022;102(3):1263–1325.

[39] Tashima R, Koga K, Sekine M, Kanehisa K, Kohro Y, Tominaga K, Matsushita K, Tozaki-Saitoh H, Fukazawa Y, Inoue K. Optogenetic activation of non-nociceptive Aβ fibers induces neuropathic pain-like sensory and emotional behaviors after nerve injury in rats. eneuro 2018;5(1).

[40] Wang H, Zylka MJ. Mrgprd-expressing polymodal nociceptive neurons innervate most known classes of substantia gelatinosa neurons. J Neurosci 2009;29(42):13202–13209.

[41] Willis WD. The role of TRPV1 receptors in pain evoked by noxious thermal and chemical stimuli. Experimental Brain Research 2009;196(1):5–11.

[42] Zhang Z, Roberson DP, Kotoda M, Boivin B, Bohnslav JP, González-Cano R, Yarmolinsky DA, Turnes BL, Wimalasena NK, Neufeld SQ, Barrett LB, Quintão NLM, Fattori V, Taub DG, Wiltschko AB, Andrews NA, Harvey CD, Datta SR, Woolf CJ. Automated preclinical detection of mechanical pain hypersensitivity and analgesia. Pain 2022;163(12):2326–2336.

